# *RecView*: an interactive R application for viewing and locating recombination positions using pedigree data

**DOI:** 10.1101/2022.12.21.521365

**Authors:** Hongkai Zhang, Bengt Hansson

**Author notes:** Author contributions: H.Z. and B.H. conceptualized the study. H.Z. developed the software with input from B.H. H.Z. performed the bioinformatic analyses. H.Z. and B.H. wrote the manuscript.

## Abstract

Recombination generates new haplotypes and disconnects linked genes thereby increasing the efficiency of selection and the adaptive potential. Quantifying the recombination landscape, *i*.*e*., the recombination rate variation along chromosomes, is important for understanding how evolutionary processes such as selection and drift are acting on genes and chromosomes. Here, we present *RecView*, an interactive R application, designed to view and locate recombination positions along chromosomes using whole-genome genotype data of a three-generation pedigree. *RecView* visualises the grandparent-of-origin of all informative alleles along each chromosome of the offspring in the pedigree. It also infers putative recombination positions with two algorithms, one based on change in the proportion of the alleles with specific grandparent-of-origin, and one on the degree of continuity of alleles with the same grandparent-of-origin, along the chromosome. Putative recombination positions are given in base pairs together with an estimated error based on the local density of informative alleles. We demonstrate the applicability of *RecView* using sequencing data of one 120 Mb-large chromosome of one offspring (and its four grandparents and two parents) of a passerine bird, the great reed warbler (*Acrocephalus arundinaceus*). On this chromosome, we detected five recombination events, three on the paternal chromosome and two on the maternal chromosome with both algorithms. To evaluate how sensitive the analysis is for SNP density, we downsampled our data to 10% of the original dataset. In conclusion, we provide an easy-to-use and highly effective application for viewing and locating recombination positions along chromosomes in small pedigrees. *RecView* and test data are available on GitHub (https://github.com/HKyleZhang/RecView.git).

## 1. Background

Recombination is an important evolutionary process because it generates novel genetic variation by reshuffling alleles and breaking the linkage between genes. Consequently, it allows linked genes to evolve partly independently and thereby makes selection more efficient. When recombination is suppressed, selection operates on large, linked regions, which lowers the fixation probability of beneficial mutations as well as the efficiency with which mildly deleterious mutations are purged. Indeed, the outcome of selection on any genomic region is dependent on the combined effect on linked loci, and the degree of linkage is in turn affected by the rate of recombination. The importance of recombination in maintaining the fitness of large genomic regions is, for example, supported by the rapid degeneration and massive gene loss on non-recombining parts of sex chromosomes (Bachtrog, 2013; Barton and Charlesworth, 1998; Charlesworth and Charlesworth, 2000; Felsenstein, 1974).

The local recombination rate covaries with several population genetic parameters, including linkage disequilibrium (LD), nucleotide diversity, GC and repeat content, and gene density (Ponnikas *et al*., 2022). These parameters are often used to infer the type and strength of selection (and other processes) acting on the genome. This warrants quantifying the recombination landscape, *i*.*e*., the local recombination rate variation along the chromosomes, in evolutionary studies. A common pattern found in many species is increasing recombination rates towards the telomeres (Haenel *et al*., 2018; Ponnikas *et al*., 2022) and exceptionally low rates around the centromeres (Limborg *et al*., 2016; Sardell and Kirkpatrick, 2020; Vincenten *et al*., 2015). However, the recombination rate often varies at a local chromosomal scale and among species, which makes such generalisations too crude and motivates gathering data on the recombination landscape of specific study species. Moreover, characterising the recombination landscape taxonomically broadly would generate data to understand other unresolved questions about recombination. For example, it is still poorly understood why the recombination rate and landscape differ between the sexes, with some species showing an absence of recombination in one sex (achiasmy) and others showing a large variation between sexes in the rate of recombination (heterochiasmy) (Burt *et al*., 1991; Haldane 1922; Huxley 1928; Lenormand 2003).

Recombination can be studied by direct observations of crossovers, or chiasmata, using cytological approaches (Hultén, 1974; Koul *et al*., 2020; Moens and Spyropoulos, 1995), or by localising proteins involved in repairing double-strand breaks (Basheva *et al*., 2008; Froenicke *et al*., 2002; Lynn *et al*., 2002). Furthermore, the recombination rate can be estimated by comparing genetic linkage maps to physical maps (Dumont *et al*., 2011; Groenen *et al*., 2009; Johnston *et al*., 2017; Robinson, 1996). Likewise, recombination rates can be estimated by quantifying the degree of LD along chromosomes, with the rationale that high recombination rates lead to faster LD decay (Mackay and Powell, 2007; Provost *et al*., 2022; Singhal *et al*., 2015). Both linkage maps and LD patterns provide estimates of the population average rate of recombination over the chromosomes. In addition, recombination positions resulting from single crossover events can be detected by investigating chromosome-wide allele sharing patterns between grandparents and grandchildren, *i*.*e*., by exploring alleles segregating in three-generation pedigrees (Smeds *et al*., 2016). The resolution of the recombination positions in such analyses is determined by the sequencing method and the degree of genetic variation of the study species, with genome-wide approaches and highly heterozygous species providing higher resolution. When such analyses are scaled to include many individuals (many meiosis), population-averaged recombination rates can also be estimated. With the recent progress in high-throughput sequencing, genome assemblies and individual genotype data can nowadays be produced for almost any study species. We believe that this development will make locating recombination positions in pedigree data increasingly common in future population genetic and evolutionary studies.

To facilitate the analysis of locating recombination positions, we present *RecView*, an interactive R application, designed to view and infer putative recombination positions along chromosomes in whole-genome sequence data in a three-generation pedigree. *RecView* requires the input of a genotype file of the individuals in a three-generation pedigree and a scaffold file providing the order and orientation of the scaffolds on each chromosome. Essentially, *RecView* analyses the grandparent-of-origin (GoO) of alleles at each single nucleotide polymorphism (SNP) in each offspring (F2 generation) separately. Specifically, the genotypes of the four grandparents, the two parents and the focal offspring of each SNP are represented by a genotype string, which is referred to as the genotype string keyword. The specific keyword for an SNP is then compared to all possible genotype string combinations in the dictionary of GoO. Some genotype combinations are informative, for example, at an SNP-locus with the two alleles (a and c), the genotypes ac_paternal grandfather_ -aa_paternal grandmother_ -aa_maternal grandfather_ -aa_maternal grandmother_ -ac_father_ -aa_mother_ -ac_offspring_ make it possible to infer the GoO of allele c in the offspring. *RecView* determines the GoO at all SNPs, and visualises the GoO of informative alleles along each chromosome (or scaffolds). Putative recombination positions can be determined by a visual inspection of the chromosome-wide GoO plot, but *RecView* also includes the option to infer putative recombination positions on the chromosomes by applying either of two algorithms: (i) the proportional difference (PD) algorithm that detects positions where the difference in the proportion of alleles with specific GoO between flanking windows reaches a local maximum, and (ii) the cumulative continuity score (CCS) algorithm that detects positions where the continuous inferences of a specific GoO switch from one grandparent to the other. The main result can be output as chromosome-wide plots (GoO-, PD- and CCS-plots), and as tables (GoO inferences at each SNP, and PD- or CCS-inferred putative recombination positions with estimated errors based on the local density of informative alleles). We demonstrate the applicability of *RecView* using short-read sequence data of one chromosome of one offspring, and its four grandparents and two parents, of a passerine bird, the great reed warbler (*Acrocephalus arundinaceus*). To evaluate how sensitive the analysis is for SNP density, we also analyse a dataset containing 10% of the original full data.

## 2. Implementation

### 2.1 Genotype data from a three-generation pedigree

*RecView* requires genotype data of bi-allelic SNPs of the individuals in a three-generation pedigree that includes four grandparents (F0), two parents (F1), and at least one offspring (F2) (**Figure 1**). The analysis is conducted independently for each offspring. The recombination being analysed occurs in the F1s, involving genetic exchange between the homologous chromosomes inherited from the F0s. If the recombined chromatid is passed down to the F2, we can trace chromosomal regions to the (paternal or maternal) grandfather or the (paternal or maternal) grandmother. The genotype file with bi-allelic SNPs can be extracted from variant calling files using, *e*.*g*., *VCFtools* (Danecek *et al*., 2011).

**Figure 1.**
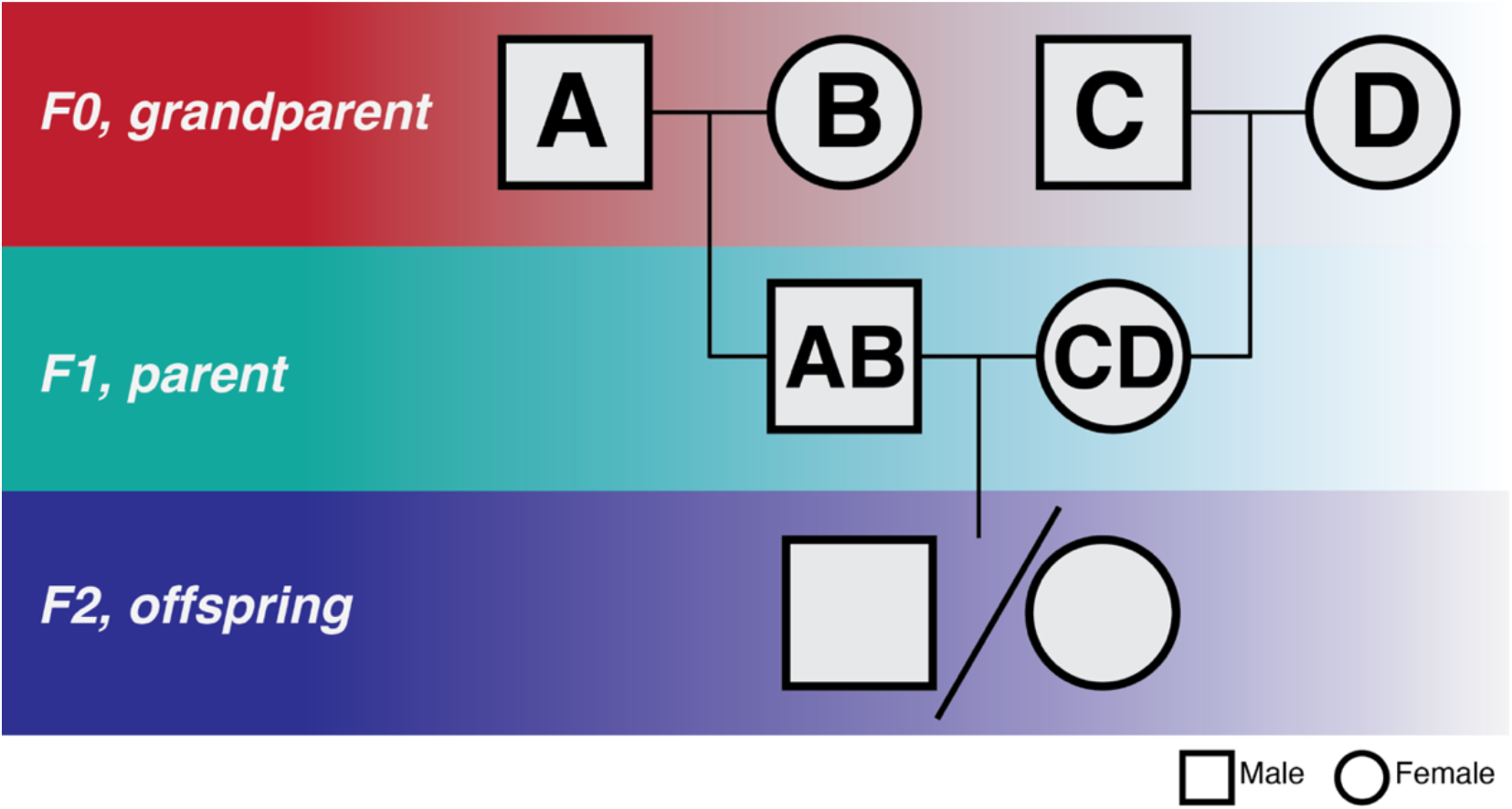
The pedigree dataset required for analysing recombination locations with *RecView*. Grandparents are labelled A, B, C and D, and parents AB and CD. The analysis is conducted independently for each offspring.

### 2.2 Dictionary of grandparent-of-origin (GoO)

#### 2.2.1 The dictionary for autosomal data

*RecView* contains a “dictionary of GoO”, which includes all possible genotype strings of seven individuals in a three-generation pedigree. Technically, each genotype at each bi-allelic SNP is represented by 0, 1 or 2 where 0 = homozygote for the reference allele, 2 = homozygote for the alternate allele, and 1 = heterozygote. This means that a genotype string for grandparents A, B, C and D, parents AB and CD, and the offspring in **Figure 1**, can be written as, *e*.*g*., 0-0-2-2-0-2-1 if both paternal grandparents are homozygous for the reference allele and both maternal grandparents are homozygous for the alternative allele, or 0-0-0-1-0-1-1 if the maternal grandmother, the mother and the offspring are heterozygous.

In total, there are 3^7^ = 2,187 different genotype strings, considering each individual has 3 possible genotypes (0, 1 or 2). Without considering mutations, we removed genotype strings that represent biologically impossible scenarios, for example, 0-0-0-0-1-0-1 where neither paternal grandparent carries the alternative allele present in the heterozygous father (and offspring). This leaves us with a dictionary of GoO with 435 biologically plausible genotype strings.

#### 2.2.2 The dictionary for sex chromosome data

A specific dictionary of GoO is constructed for the X and Z chromosomes, or more precisely for the part of these chromosomes where the heterogametic sex does not recombine (XY males and ZW females). Basically, the dictionary of GoO is modified so that the GoO is not inferred for the paternal X chromosome in XY system and the maternal Z chromosome in ZW system.

#### 2.2.3 The usage of the dictionary

As the dictionary of GoO contains all possible genotype strings and their corresponding inferred GoO, the 7-individual genotype string of each SNP can be used as keywords in the dictionary to obtain the GoO inference. The result can be visualised as the GoO of each SNP ordered by positions. **Figure 2** shows a hypothetical example of GoO of 200 SNPs along a small chromosome, which is also used for demonstrating the two algorithms to automatically locate recombination. *RecView* does not plot unresolved and thus uninformative alleles. Note that in this example, we use the order of the SNPs as their positions, which is not the case for real data where the actual positions on the chromosome are used for plotting and for interpretation of the actual recombination positions in the unit of base pair.

**Figure 2.**
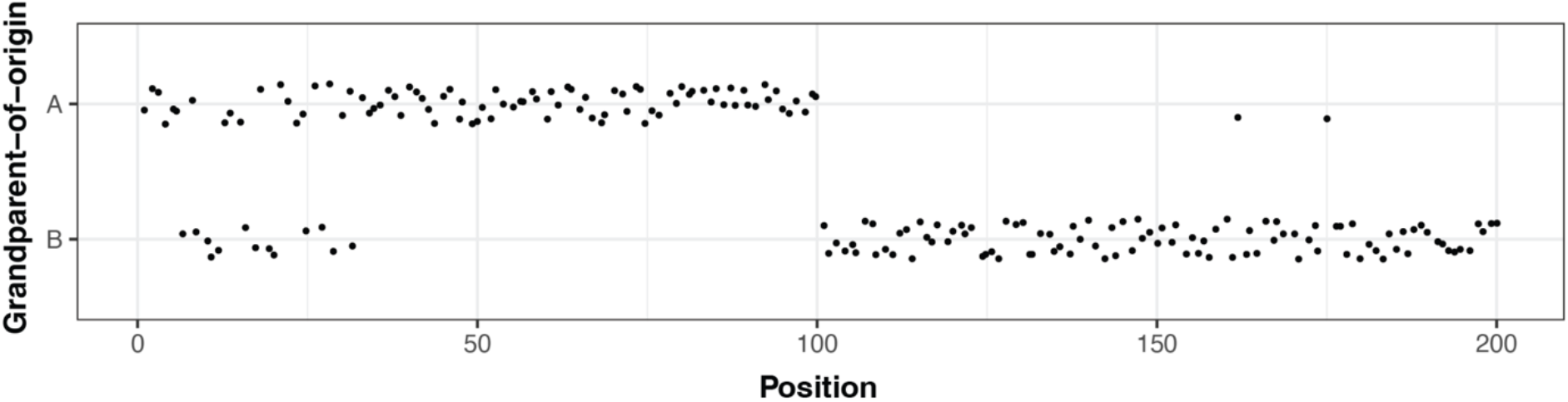
The paternal grandparent-of-origin inference for 200 SNPs. These data are hypothetical but were selected to indicate how, *e*.*g*., sequencing errors may affect patterns. Data points are assigned with noise on the y-axis to avoid overlapping.

### 2.3 Locating recombination positions

#### 2.3.1 Proportional difference (PD) algorithm

The PD algorithm calculates the absolute difference in the proportion of alleles originating from a specific grandparent between flanking windows. It proceeds by specifying a window size (the number of informative SNPs of each flanking window), a step value (k) giving the number of SNPs between each calculated position, and a threshold to trigger denser calculations (*e*.*g*., at every SNP) to detect local maxima.

Taking the example of the paternal chromosome, the evaluation at each position starts with calculating the proportion of alleles originating from grandparent A for the flanking windows (S1 for the downstream and S2 for upstream). Next, the absolute difference of the proportion A for S1 and S2 (| Δ_*GoO prop*_.|) is calculated for every k bp. If | Δ_*GoO prop*_.| is smaller than the user-specified threshold, the process proceeds every k SNP, but if it is equal to or larger than the user-specified threshold, | Δ_*GoO prop*_.| be calculated for every informative SNP starting from the previous focal position (*i*.*e*., current position - k). The step will return to a step size of k when | Δ_*GoO prop*_.| is again smaller than the threshold. This design is intended to increase calculation speed while keeping the accuracy of the analysis. Putative recombination positions are defined as local maxima in | Δ_*GoO prop*_.| the threshold. **Figure 3** depicts the result of a PD analysis of the 200 SNPs in **Figure 2** with settings of k = 5, window size = 19, and threshold = 0.85. The process of taking finer step starts from position 95 because position 100 is the first position (when using k = 5) with | Δ_*GoO prop*_.| ≥ 0.85.

**Figure 3.**
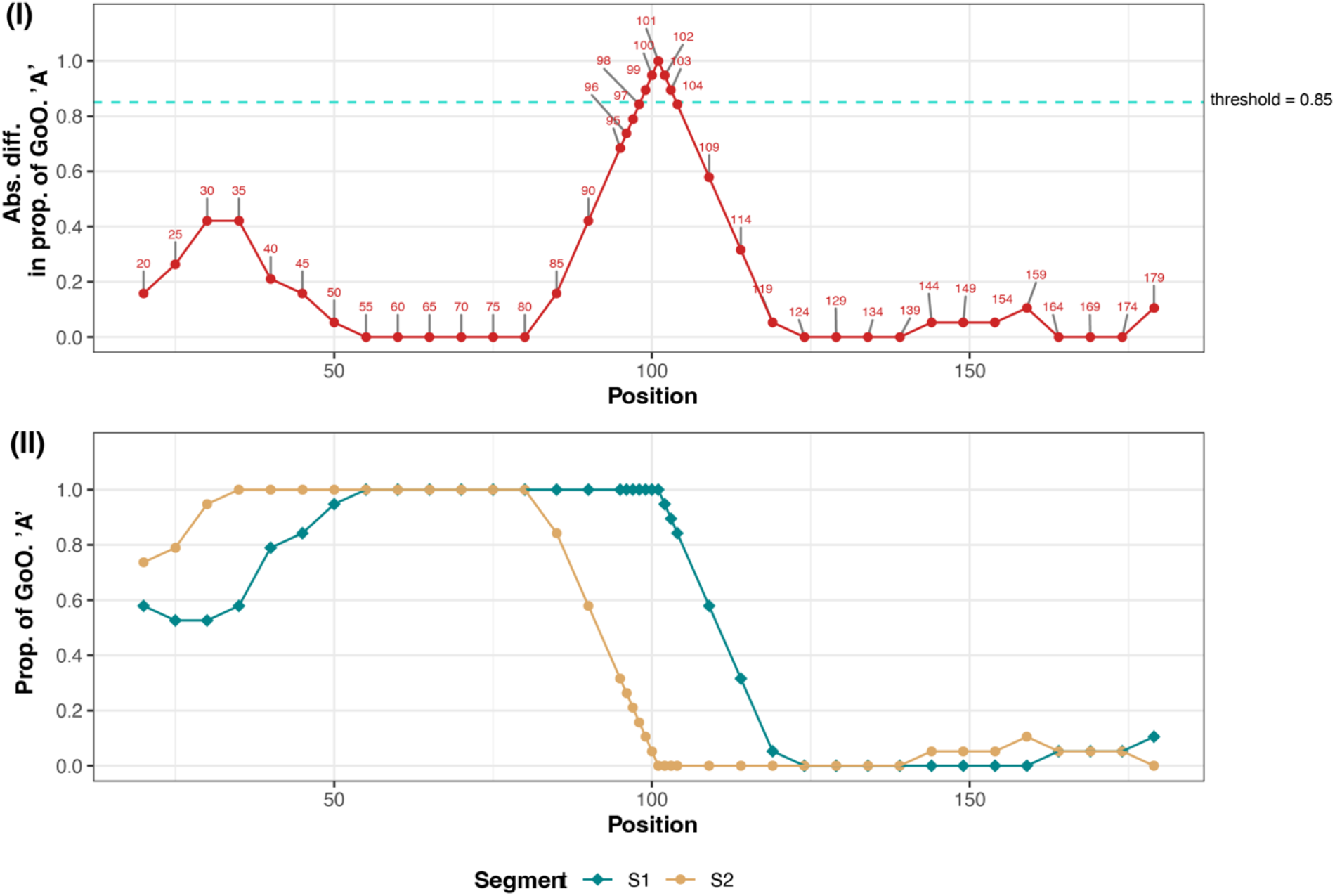
Illustrative demonstration of the sliding window algorithm to automatically locate recombination positions. (I) The absolute difference of the proportion of the grandpaternal allele A for downstream (S1) and upstream (S2) windows (| Δ_*GoO prop*_.|) along the chromosome. (II) The proportion of the grandpaternal allele A for S1 (green) and S2 (yellow), respectively, at each focal position. Position 101 shows a local maximum above the threshold and is thus a putative recombination position.

#### 2.3.2 Cumulative continuity score (CCS) algorithm

The CCS algorithm calculates a CCS for each position along the chromosome. The CCS describes the number of consecutively proceeding GoO inferences being the same as for the focal position (*e*.*g*., CCS = 3 for the last A in BAAAA). The algorithm finds putative recombination positions by locating regions where long continuously increasing slopes of CCSs of one GoO is replaced by long continuously increasing slopes of CCSs from the other grandparent.

Taking the example of the paternal chromosome, if both the focal and the previous position have GoO inference A, the focal position gets a continuity score of +1. Similarly, if the GoO inference is B for both the focal and the previous position, the focal position gets a continuity score of −1. In contrast, if the focal and the previous positions have different GoO inferences, the focal position will reset the CCS to 0. Plotting the CCS of each position along a chromosome will create increasingly positive and increasingly negative slopes (**Figure 4**) in which positive slopes indicate that the informative alleles in this region originate from the grandfather, whereas negative slopes indicate alleles originating from the grandmother.

**Figure 4.**
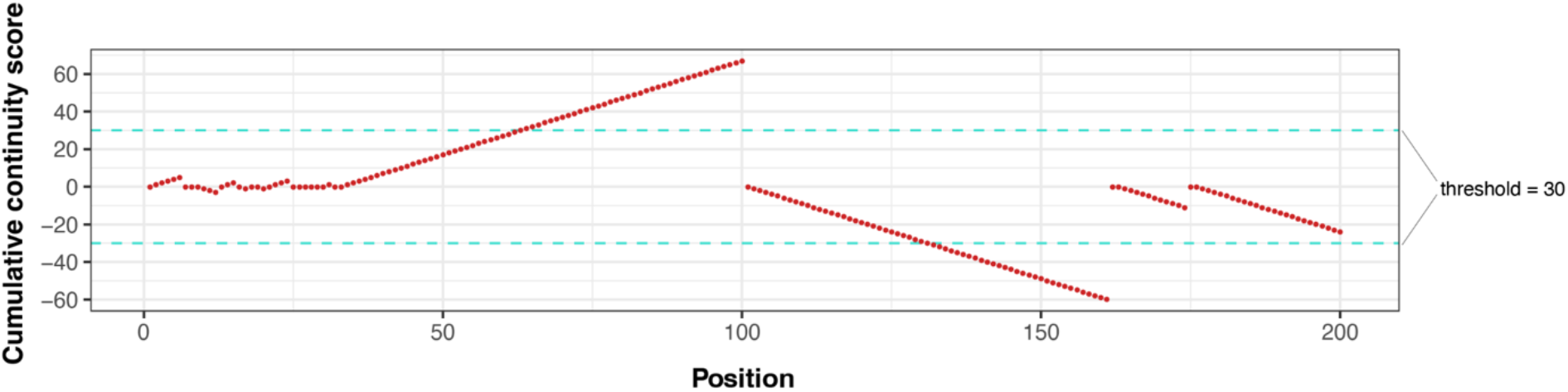
Illustrative demonstration of the cumulative continuity score (CCS) algorithm to automatically locate putative recombination positions. The CCS is reset to zero whenever the next grandparent-of-origin (GoO) inference is different. Positive and negative CCSs indicate the continuity of GoO A and B, respectively. With a threshold of CCS = 30, there is one putative recombination position, between positions 100 and 101.

To locate recombination positions, a user-specified threshold is used to exclude noise from mapping errors and wrongly called genotypes (which will reset CCS to 0). The selected threshold will depend on assumptions of the minimal size of the recombined regions, the density of the informative alleles, and the amount of noise in the data. **Figure 4** depicts the output of the 200 SNPs in **Figure 2** with a CCS threshold = 30, which implies that putative recombination positions will be located between the end of a positive (or negative) slope reaching above CCS = 30 and the beginning of the next negative (or positive) slope reaching above CCS = 30. We refer to these these two positions as the left and right observed boundaries. In addition, note that slopes with values below the threshold might occur between these slopes (which is not the case in this example). In all cases, the putative recombination positions will be assigned the middle position between the left and right observed boundaries. In the example shown in **Figure 4**, a putative recombination position occurs between positions 100 and 101.

#### 2.3.4 Uncertainty of putative recombination positions

To estimate the uncertainty or precision of putative recombination positions, we calculate the local density of informative alleles in non-overlapping sliding windows of 100 Kb. The estimated error is calculated by the reverse local density of informative alleles (*i*.*e*., 1/local density of informative alleles), *i*.*e*., the average number of base pairs per informative allele (unit: bp/allele). Putative recombination positions located in a region with low local density have higher estimated error and lower certainty than those located in a region with high local density.

### 2.4 *RecView* interactive R application

We implement the GoO inference, and the PD and CCS algorithms, into a ShinyApp wrapped in the R package *RecView*. The *RecView* is intended to provide an easy-to-use GUI to view and locate recombination positions using pedigree data. The basic workflow of *RecView* is shown in **Figure 5**.

**Figure 5.**
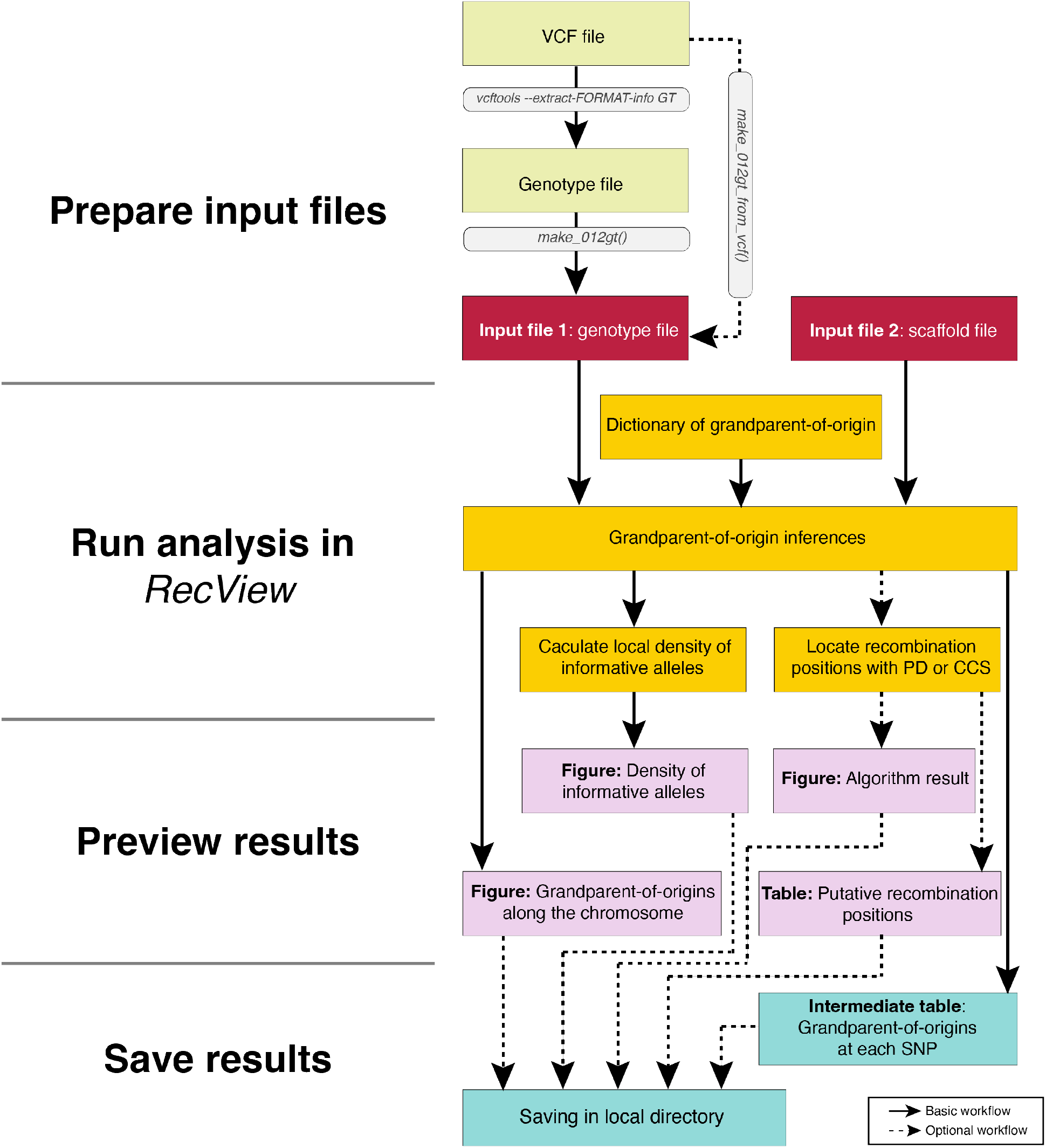
The workflow of using *RecView*. Solid lines indicate the basic workflow while dashed lines indicate the optional workflow. *RecView* requires an input genotype file which can be generated by using *make_012gt()* on the output file from *VCFtools*, or using *make_012gt_from_vcf()* on the VCF file. *RecView* further requires an input scaffold file containing the order and orientation of the scaffolds. These two input files are used together with the built-in dictionary of grandparent-of-origin (GoO) to produce the GoO figure showing the GoO inferences of alleles along the scaffolds, and a figure showing the informative alleles density. *RecView* can further locate putative recombination positions with the proportional difference or cumulative continuity score algorithms and output result figures and tables. The result figures and tables can be saved, including an intermediate table containing the GoO inferences at each SNP.

#### 2.4.1 Input files

As inputs, *RecView* requires a genotype and a scaffold file. *RecView* provides two functions to help generate the correct genotype file:

- *make_012gt()*: Generates the required 012-formatted genotype file from an output file constructed by using the option “--extract-FORMAT-info GT” in *VCFtools*.
- *make_012gt_from_vcf()*: Generates the required 012-formatted genotype file from a VCF file.

The scaffold file, which provides the order and orientation of the scaffolds, needs to have five columns with the following headings (case sensitive): “scaffold” (the label of the scaffold; character), “size” (the size of the scaffold in bp; integer), “CHR” (the chromosome the scaffold belongs to; character), “order” (the order of the scaffold on the chromosome; integer), and “orientation” (the scaffold orientation on the chromosome; + or -). Data for each scaffold are given in separate rows.

#### 2.4.2. Using *RecView*

*RecView* is executed by the command *run_RecView_App()* (**Figure 6**) in Rstudio (Racine, 2012). It uses the local machine as the server, which means that it can be run offline. In the GUI, the user can upload the two required input files and select settings, including which chromosome(s) and individuals to visualise, whether putative recombination positions should be calculated and if so with which algorithm (PD or CSS), whether or not the result figures and/or tables should be saved locally, *etc*. The output panel shows result figures and tables in different tabs (**Figure 6**).

**Figure 6.**
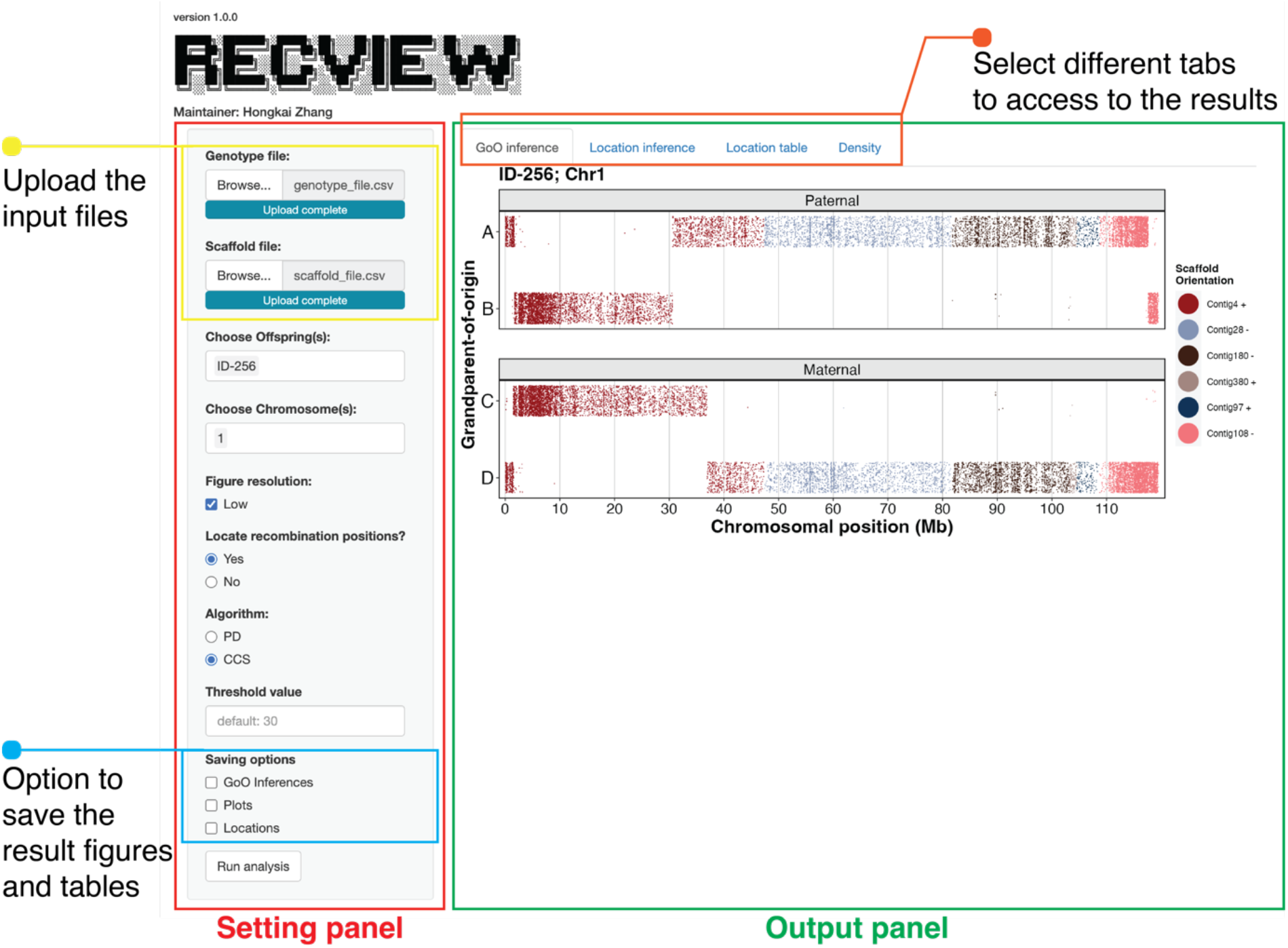
The GUI of *RecView* with the setting panel (red square) for uploading input files (yellow square) and setting options and the output panel (green square) where results can be accessed by selecting different tabs (orange sqaure).

## 3. Results

### 3.1 Example dataset from a three-generation pedigree of great reed warbler

We demonstrate the usage of *RecView* with short-read sequencing data from a three-generation pedigree of a passerine bird, the great reed warbler. The sequencing data contain 150-bp paired reads with 350-bp insert size at approximately 50X depth coverage. The reads were trimmed and mapped to the great reed warbler reference genome (Sigeman *et al*., 2021), and genotypes of the seven individuals (four grandparents, two parents and an offspring called ID-256) were called with *freebayes* (Garrison and Marth, 2012) and saved as a VCF file. We kept bi-allelic SNPs and used *VCFtools* (option: --extract-FORMAT-info GT) to extract the genotypes from the VCF file. We formatted the input genotype file using the *make_012gt()* function. In addition, we downsampled the number of SNPs to 10% of the original number (referred to as the “downsampled dataset”), to mimic the situation where fewer SNPs are available for the analysis, such as for reduced representation sequencing data (*e*.*g*., restriction site-associated (RAD) sequencing data; *e*.*g*., Hansson *et al*., 2018). For the scaffold file, we used ordered and oriented scaffolds of the great reed warbler assembly (Sigeman *et al*., 2021; B. Hansson *et al*., unpubl.). As a test data set, data for two offspring (ID-256, ID-258) and their grandparents and parents at two chromosomes (chromosome 1 and 28) are provided (available on *RecView*’s GitHub page).

To locate putative recombination positions, we selected a window size of 550 SNPs, a step of k = 17 SNPs, a finer step of 1 SNP, and threshold of 0.9, for the proportional difference (PD) algorithm, and a threshold of 30 for the cumulative continuity score (CCS) algorithm.

### 3.2 Grandparent-of-origin (GoO) inference plots

Figure 7 shows the informative alleles of SNPs along chromosome 1 in offspring ID-256 on the paternal and maternal sides as inferred by *RecView* for the full and downsampled datasets, respectively. The data of SNPs on different scaffolds are plotted in different colours. Putative recombination positions can be visually located and in this case, we suggest three crossovers on the paternal chromosome and two on the maternal chromosome in both the full and reduced data (**Figure 7**).

**Figure 7.**
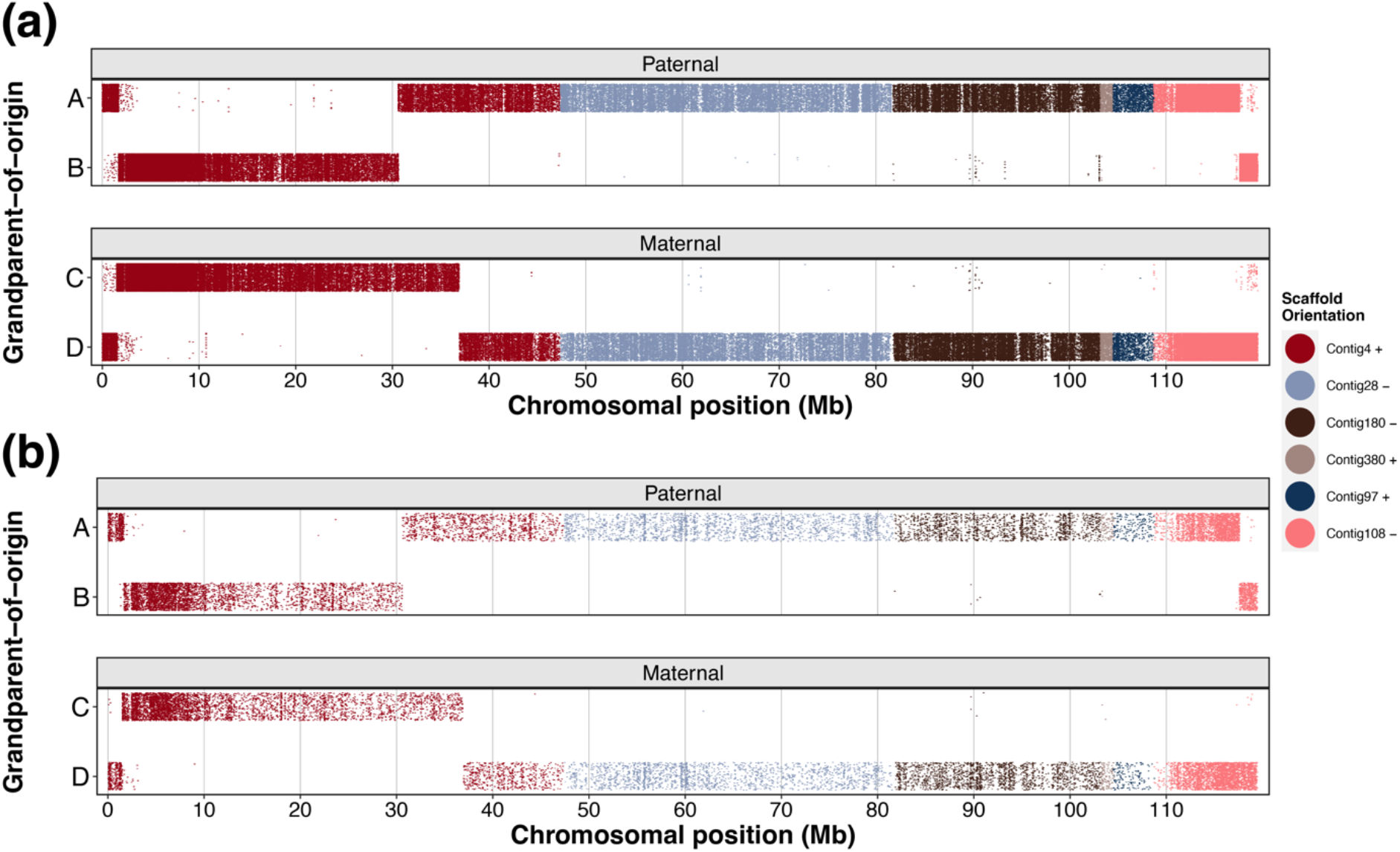
The grandparental-of-origin of informative alleles at all SNPs along chromosome 1 in great reed warbler offspring ID-256 for (a) the full dataset and (b) the downsampled dataset. Each dot represents an allele at a specific SNP for the paternal or maternal chromosomes. Dots are plotted with noise on the y-axis to alleviate the degree of overlap. Colouration indicates different scaffolds on chromosome 1 in the great reed warbler genome assembly (Sigeman *et al*., 2021).

### 3.3 Putative recombination positions inferred from the PD algorithm

With the selected threshold of 0.9, the PD algorithm suggests five recombination events, three in the paternal chromosome and two on the maternal chromosome, and **Figure 8** shows the output plot from *RecView*. The local density of the informative allele, which is used as an estimated error of the putative recombination positions, is also calculated and plotted (**Figure 9**). The recombination positions and estimated errors are outlined in **Table 1**, where it can be observed that the resolution of most recombination positions is < 1kb. Compared to the full dataset, the downsampled dataset resulted in decreased resolution with approximately 10-15 times larger estimated errors, and two recombination positions near the chromosome ends were missed (**Figure 9, Table 1**).

**Table 1.**
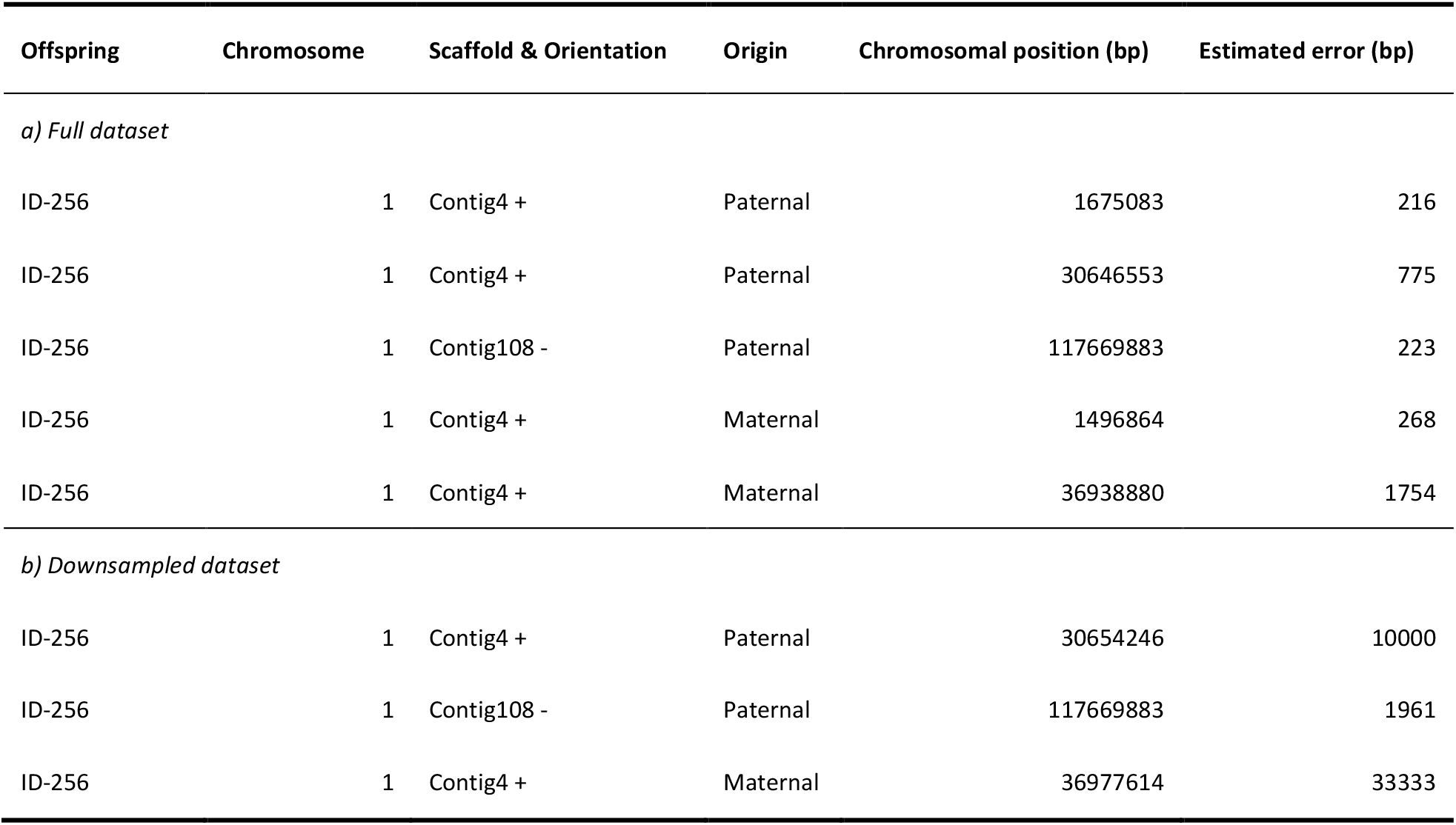
Putative recombination positions and estimated errors inferred from the proportional difference algorithm in the full dataset (a) and downsampled dataset (b).

| Offspring | Chromosome | Scaffold & Orientation | Origin | Chromosomal position (bp) | Estimated error (bp) |
| --- | --- | --- | --- | --- | --- |
| *a) Full dataset* |  |  |  |  |  |
| ID-256 | 1 | Contig4 + | Paternal | 1675083 | 216 |
| ID-256 | 1 | Contig4 + | Paternal | 30646553 | 775 |
| ID-256 | 1 | Contig108 - | Paternal | 117669883 | 223 |
| ID-256 | 1 | Contig4 + | Maternal | 1496864 | 268 |
| ID-256 | 1 | Contig4 + | Maternal | 36938880 | 1754 |
| *b) Downsampled dataset* |  |  |  |  |  |
| ID-256 | 1 | Contig4 + | Paternal | 30654246 | 10000 |
| ID-256 | 1 | Contig108 - | Paternal | 117669883 | 1961 |
| ID-256 | 1 | Contig4 + | Maternal | 36977614 | 33333 |

**Figure 8.**
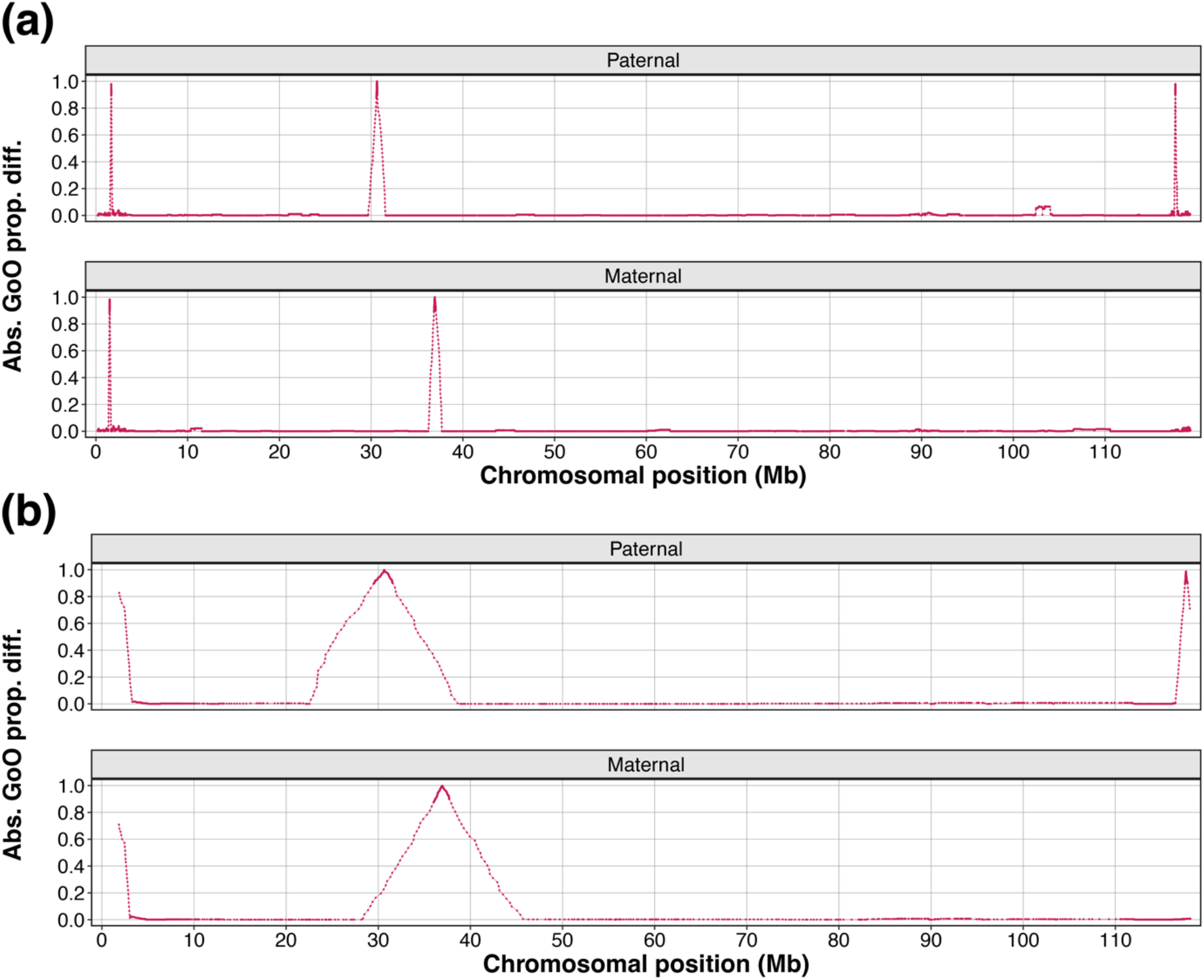
Visualization of the result from the proportional difference algorithm to locate putative recombination positions. Shown are the absolute difference of the proportion of the grandpaternal allele A, and C for downstream (S1) and upstream (S2) windows (| Δ_*GoO prop*_.|) along the Chromosome 1 in offspring ID-256 in the full dataset (a) and downsampled dataset (b).

**Figure 9.**
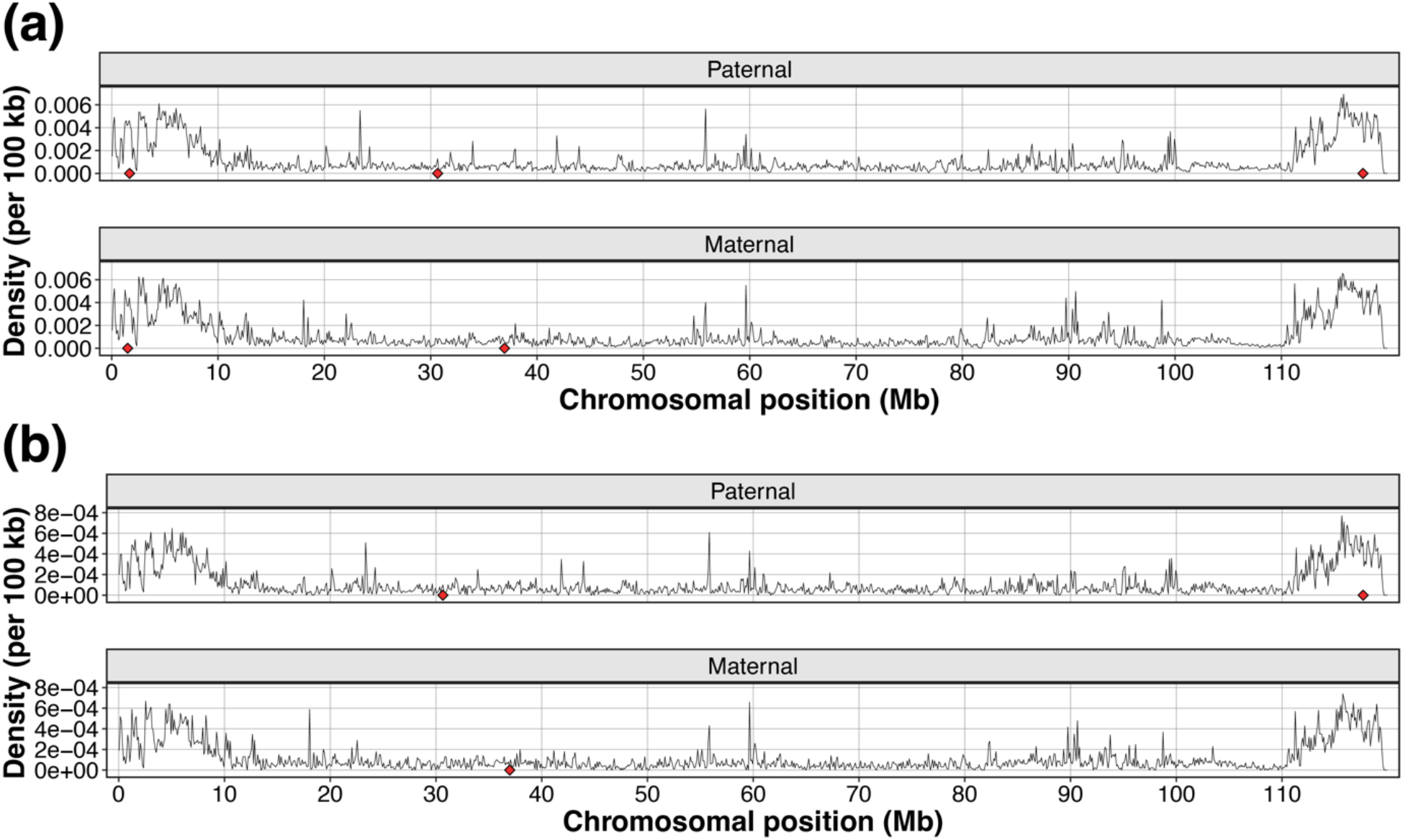
The local density of informative SNPs along the Chromosome 1 in offspring ID-256 in the full dataset (a) and downsampled dataset (b). The red diamonds show the putative recombination positions from the proportional difference algorithm.

### 3.4 Putative recombination positions inferred from the CCS algorithm

With the selected threshold of 30, the CCS algorithm suggests five recombination events, three in the paternal chromosome and two on the maternal chromosome, and **Figure 10** shows the output plot from *RecView*. As for the PD algorithm, the local density of the informative allele, which is used as an estimated error of the putative recombination positions, is also plotted (**Figure 11**). The recombination positions and estimated errors are outlined in **Table 2**. Compared to the full dataset, the downsampled dataset resulted in decreased resolution (larger errors), but still all five putative recombination positions were located (**Figure 11, Table 2**).

**Table 2.** Putative recombination positions inferred from the cumulative continuity score algorithm in the full dataset (a) and the downsampled dataset (b). Also given are estimated error, and left and right observed boundaries.

| Offspring | Chromosome | Scaffold & Orientation | Origin | Chromosomal position (bp) | Estimated error (bp) | Left observed boundary (bp) | Right observed boundary (bp) |
| --- | --- | --- | --- | --- | --- | --- | --- |
| *a) Full dataset* |  |  |  |  |  |  |  |
| ID-256 | 1 | Contig4 + | Paternal | 1674925 | 216 | 1674767 | 1675083 |
| ID-256 | 1 | Contig4 + | Paternal | 30641720 | 775 | 30636886 | 30646553 |
| ID-256 | 1 | Contig108 - | Paternal | 117669374 | 223 | 117668866 | 117669883 |
| ID-256 | 1 | Contig4 + | Maternal | 1496182 | 268 | 1495501 | 1496864 |
| ID-256 | 1 | Contig4 + | Maternal | 36936350 | 1754 | 36933819 | 36938880 |
| *b) Downsampled dataset* |  |  |  |  |  |  |  |
| ID-256 | 1 | Contig4 + | Paternal | 1674915 | 2564 | 1674316 | 1675514 |
| ID-256 | 1 | Contig4 + | Paternal | 30635440 | 10000 | 30616635 | 30654246 |
| ID-256 | 1 | Contig108 - | Paternal | 117668860 | 1961 | 117667836 | 117669883 |
| ID-256 | 1 | Contig4 + | Maternal | 1497034 | 3125 | 1495311 | 1498757 |
| ID-256 | 1 | Contig4 + | Maternal | 36939704 | 33333 | 36901793 | 36977614 |

**Figure 10.**
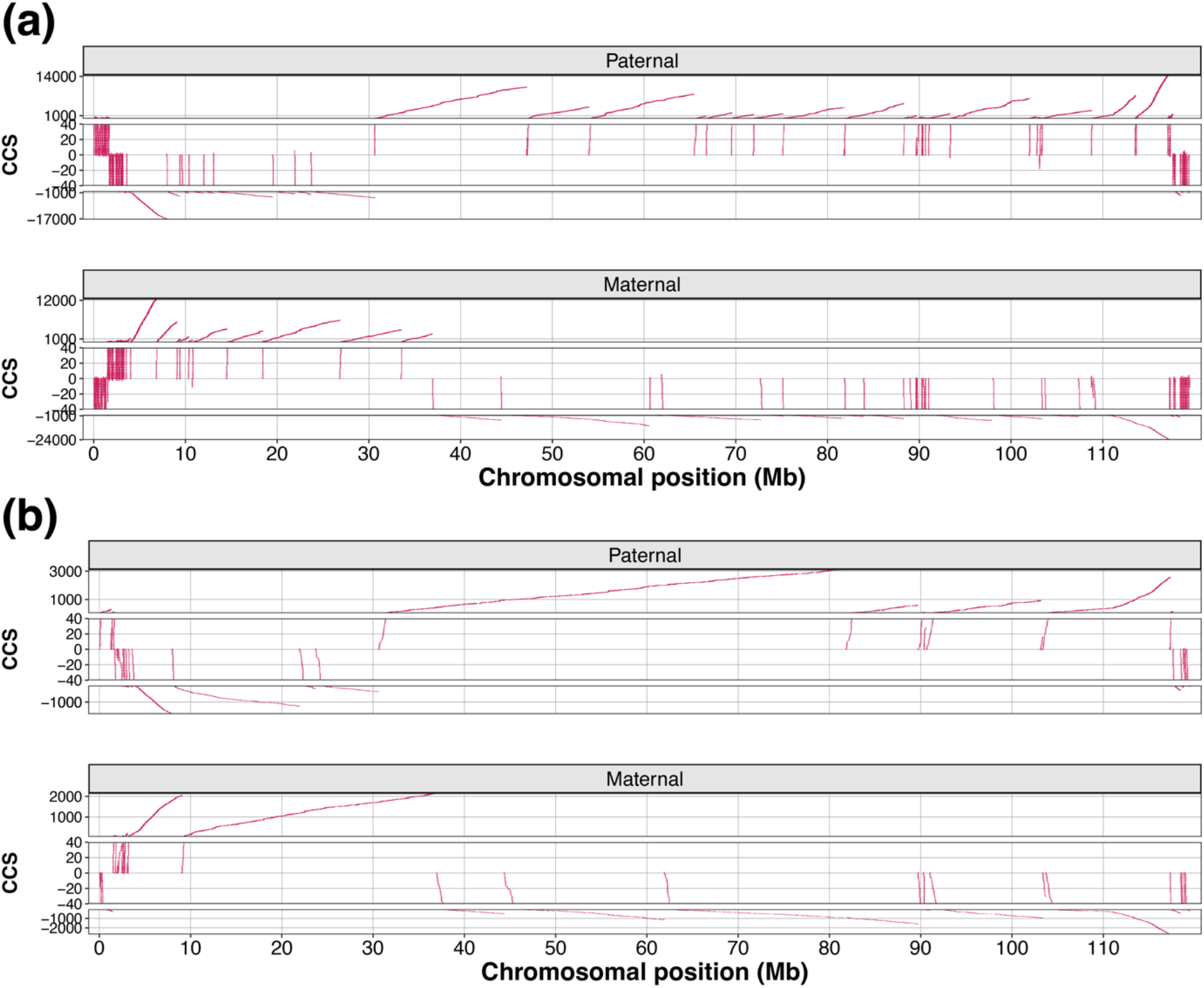
Visualization of the result from the cumulative continuity score (CCS) algorithm to locate putative recombination positions. Shown are the CCS for the maternal and maternal chromosomes along the Chromosome 1 in offspring ID-256 in the full dataset (a) and the downsampled dataset (b). Increasing slopes indicate continuous alleles with inferred GoO from the paternal grandfather (upper panel) or maternal grandfather (lower panel), while decreasing slopes indicate the origin of the paternal grandmother (upper panel) or maternal grandmother (lower panel).

**Figure 11.**
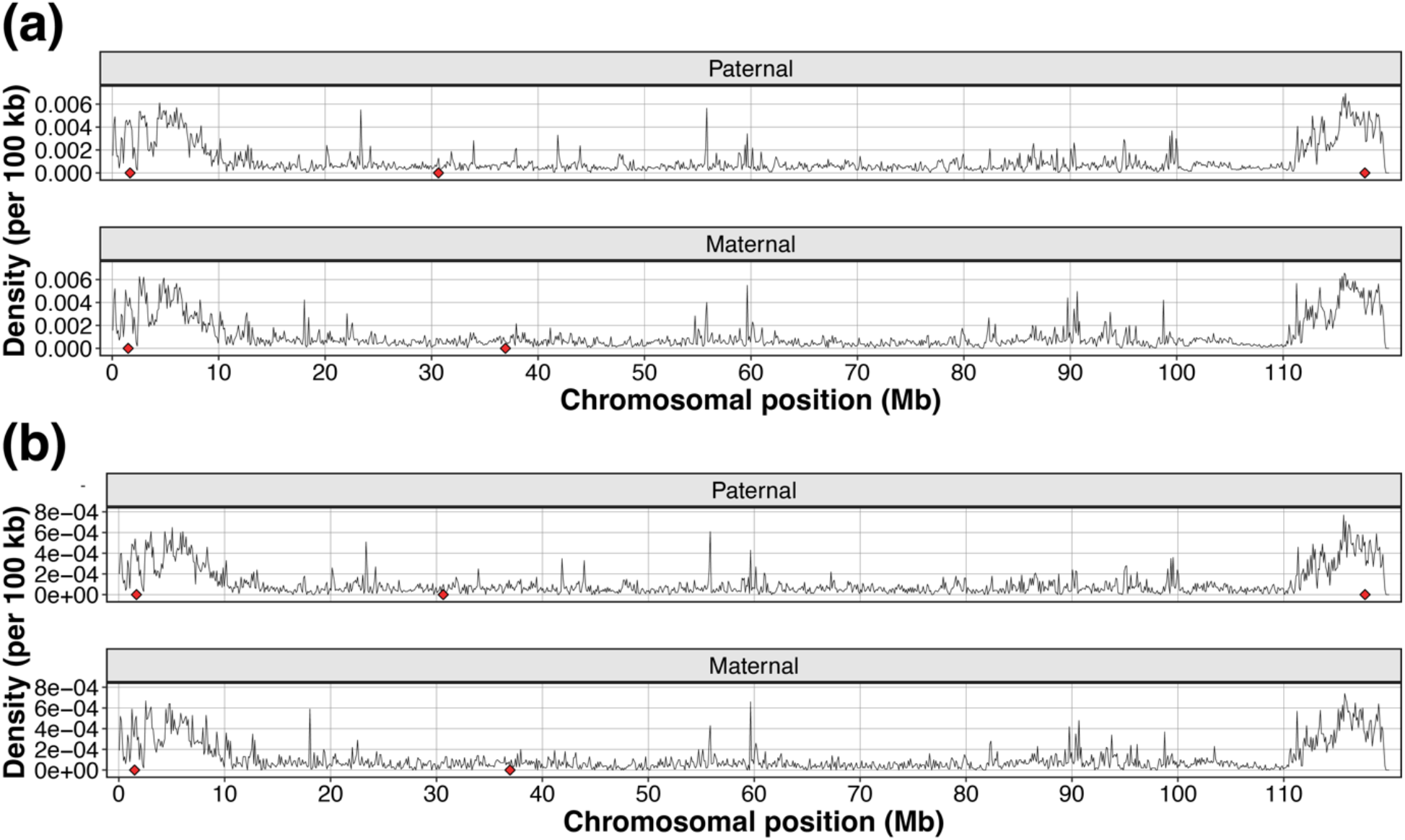
The local density of informative SNPs along chromosome 1 in offspring ID-256 in the full dataset (a) and downsampled dataset (b). The red diamonds show the putative recombination positions from the cumulative continuity score algorithm.

## 4. Discussion

The recombination landscape with regions of high and low recombination rates can be quantified by estimating genetic distances between markers segregating in large pedigrees (Groenen *et al*., 2009; Johnston *et al*., 2017) and by using population-level estimates of LD (Provost *et al*., 2022; Singhal *et al*., 2015). Another approach to study recombination is to locate actual recombination positions on individual chromosomes originating from single meiotic events (Bell *et al*., 2020; Smeds *et al*., 2016). This can be done, as in the present study, by the principle to find the positions in the genome that separate adjacent chromosomal regions with different grandpaternal and grandmaternal origins, respectively.

We present and demonstrate *RecView*, an interactive R application, which facilitates viewing and locating recombination positions using high-throughput sequencing data of three-generation pedigrees. *RecView* provides an interactive GUI for easy run of analyses, choosing different parameter settings, previewing and saving results. *RecView* infers grandparent-of-origin (GoO) for the alleles at each SNP efficiently by utilising the genotypes of all analysed individuals in the pedigree to construct a genotype string that then is used to infer the origin of each allele by comparing to a dictionary of GoO. This process of matching genotypes and obtaining information of GoO is computationally faster than executing a series of conditional processes, because the same inferring processes given the same genotype combination will not be redundantly conducted and also the GoO inference given a specific genotype string is already conducted during the *a priori* construction of the dictionary.

*RecView* analyses all genotypes provided in the input file, including possibly wrongly called genotypes caused by sequencing and mapping errors. Some wrongly called SNPs will result in impossible genotype strings and will not be included in the output, whereas others may result in noise in the data, causing contrasting GoO inferences compared to adjacent SNPs along the chromosome (**Figure 7** where several SNPs, *e*.*g*., in the first 3 Mb-region, appear to be genotype errors). Crossovers are commonly separated by large chromosome regions which means that some level of noise in the data is acceptable; the large-scale patterns of chromosomal regions segregating in the pedigree will be detected anyway. However, naturally, the larger the noise in relation to the SNP density, the more problematic inferring crossovers will become. Thus, erroneously called genotypes can confound detecting real recombination events, especially when the size of the recombined region is small. This is rarely the case for crossovers as recombination interference separates crossover events (Coop and Przeworski, 2007), but could be a serious problem if the aim is to infer gene conversion events (non-crossovers) which typically span merely a few hundred base pairs (Duret and Galtier, 2009; Gergelits *et al*., 2021). Noise in the data can be reduced by deeper sequencing and/or stricter filtering criteria during the SNP calling phase of the project, *i*.*e*., steps before the *RecView* analyses.

*RecView* implements two algorithms to locate recombination positions, the proportional difference (PD) or cumulative continuity score (CCS) algorithms. The PD algorithm finds positions where the two flanking windows differ as much as possible in which grandparental alleles they capture. The window size is selected by the user and sets a limit for how small regions can be detected and how close to the chromosome ends recombination events can be detected. This restriction may be partly avoided by using different window sizes in the PD analysis. CCS finds positions between two regions having a sufficient number of continuous informative alleles from different grandparents. The CCS algorithm may be able to locate recombination events also close to the chromosome ends, but may in general be more sensitive to wrongly called genotypes than the PD algorithm, because the CCS of a stretch of informative alleles from the same grandparents will be disrupted by errors (set to 0) and thus stay below the specified threshold, which means that large CCS can be missed (*e*.*g*., region 0-3 Mb in **Figure 7a** and **10a** where frequent noise causes relatively short CCSs). Due to these pros and cons of the two algorithms, we recommend using both when studying recombination events, in particular for species where recombination is strongly biased towards the telomeric ends of chromosomes.

The resolution of the inferred recombination positions is dependent on the size of the recombined region and the distribution of informative SNPs. In regions with dense informative SNPs, the inferred recombination positions are more likely located close to informative alleles and thus have high resolution. This means that there is a negative association between the SNP density and the resolution of recombination positions, which is why we provide the reverse local density of informative alleles as an estimated error for putative recombination positions. The reverse local density of informative alleles indicates the size (in bp) that an informative SNP covers, and differs between species (due to differences in heterozygosity) and sequencing techniques (due to differences in the number of called SNPs).

It is important to consider the completeness of the genome assembly when analysing recombination, because crossovers in unassembled parts of genomes will be missed. We recommend careful consideration of each analysed chromosome arm which should have at least a 50 cM recombination distance due to a requirement of one obligate crossover (Stapley *et al*., 2017). Thus, chromosome arms with atypically few detected recombination events may represent incompletely assembled genome regions. Likewise, wrongly assembled chromosomes may result in erroneous inferences of recombination numbers and positions. For example, if Contig 108 were to be wrongly oriented in our great reed warbler assembly (+ instead of -), another crossover event would have been identified and thus led to a quite small double-crossover event in the parental chromosome (**Figure 7**). An unintended application of *RecView*, when data of several meiotic events are available, is to curate genome assemblies. In situations where the analyses of multiple offspring result in repeated putative recombination positions at the same, specific position in multiple offspring (and in particular if this coincides with scaffolds boundaries) one may suspect assembly errors which may be considered to be corrected.

## 5. Conclusions

*RecView* provides an easy-to-use GUI to facilitate viewing and locating recombination positions using genome-wide data in a three-generation pedigree. We applied *RecView* on a great reed warbler pedigree to demonstrate the features including visualisation of informative alleles with grandparent-of-origin (GoO) at SNPs along a chromosome, which makes visually detecting putative recombination positions possible. We further show how putative recombination positions can be located by two algorithms implemented in *RecView*; the proportional difference (PD) and cumulative continuity score (CCS) algorithms. The algorithms have their strengths and weaknesses, and we recommended using both to locate recombination positions.

## Notes

Conflict of Interest: The authors have no conflict of interest to declare.

### Competing Interest Statement

The authors have declared no competing interest.

